# Long-lasting negative effects of past forestation on grassland pollination networks and their function

**DOI:** 10.1101/2024.07.02.601666

**Authors:** Gaku S. Hirayama, Taiki Inoue, Tanaka Kenta, Hiroshi S. Ishii, Atushi Ushimaru

**Affiliations:** Graduate School of Human Development and Environment, Kobe University, 3-1, 1Tsurukabuto, Kobe 657-8501, Japan

**Keywords:** grassland biodiversity, ski grassland, legacy effect, plant-pollinator network generalization/specialization, pollen flow, pollen limitation, long-term management

## Abstract

1. Grasslands are facing a rapid decline worldwide. Among the primary threats to these ecosystems are changes in land use, such as land abandonment and forestation, which promote forest vegetation in detriment of grassland plant diversity. To support the conservation and restoration of grasslands, it is key to understand what ecological processes limit the recovery of their biodiversity and functions after perturbations. However, we still know little about the legacy effects of forestation on the ecological mechanisms involved in the recovery of grasslands, especially concerning long-lasting impacts on plant-pollinator interaction networks and plant reproduction.
2. Here, we aim to fill this gap of knowledge by comparing the plant and pollinator diversity, the degree of network generalization, the pollination success and pollen limitation of native plant species in 30 plant-pollinator networks of old-growth and restored grasslands of different ages since recovery (from 2 to 84 years). We hypothesized that past forestation would exert long-lasting legacy negative effects on plant richness and plant-pollinator networks, increasing pollen limitation for native plants, thereby delaying community recovery in restored grasslands.
3. Results showed that restored grasslands exhibited significantly lower plant richness, less specialized (more generalized) interaction networks, lower pollination success and pollen-limited seed reproduction of native plants compared to old-growth grasslands. Meanwhile, the degree of network specialization and pollination success gradually increased with time after grassland restoration initiated. Overall, network generalization, which was caused by low plant richness, reduced pollination and reproduction success in native grassland plants, and degraded pollination networks and functions could recover in restored grasslands with continuous management. These findings imply that plant diversity restoration was slow because of the negative feedbacks associated with low plant richness and consequently, generalized plant-pollinator interaction networks, which diminished native plant reproduction in restored grasslands.
4. *Synthesis and applications* Our findings suggest that the recovery of specialized plant-pollinator networks by enhancing plant diversity is essential for restoring pollination function. For quicker grassland restoration, it may be effective to facilitate the establishment of highly specialized pollination networks by seeding or planting diverse native plants collected from neighbouring areas while avoiding genetic contamination.

## INTRODUCTION

Grasslands are among the largest terrestrial biomes worldwide (White et al., 2000) and have historically increased over thousands of years during the Holocene through human land management such as burning, livestock grazing, and mowing (San Miguel et al., 2016). Semi-natural grasslands have declined globally over the past century due to conversion to arable lands, land abandonment resulting in woody vegetation, and afforestation, all of which cause a serious loss of grassland biodiversity (Partel et al., 2005; Torok & Dengler, 2018). To counteract these rapid losses of grassland biodiversity and their ecosystem services, restoring abandoned grasslands by removing woody vegetation and resuming management are major and urgent challenges (Pykala, 2003; Helm et al., 2019). Despite continuous restoration efforts conducted over several decades, the diversity of native plant and animal species is often lower in restored grassland communities compared to old-growth grasslands (Buisson et al., 2018; Coffey et al., 2018; Helm et al., 2019).

Recent studies have shown that past forestation can have long-lasting legacy effects on plant diversity in restored grasslands. These legacy effects have been attributed to the alterations in the soil physiochemical properties (Del Fabbro and Prati, 2015). Previous studies have also focused on the negative legacy effects on plant communities caused by the loss or absence of soil seed banks (Stahlheber et al., 2015), seed dispersal limitation (Myers & Harms, 2009; Buisson et al., 2019), and modified abiotic and biotic interactions among plants, soil properties, and microbes (Stover & Henry, 2019; Wolfsdorf et al., 2021). A recent study revealed that particularly insect-pollinated plant species take longer to recover after grassland restoration compared to wind-pollinated plants (Albert et al., 2021). This suggests that the negative legacy effects of forestation on restored grasslands can also be due to degraded plant-pollinator mutualistic interactions and the resulting reduction in reproductive success in plants. Yet, this aspect has been little explored to date. Nearly 80% of flowering plants depend on wild pollinators for their reproduction in temperate regions (Ollerton et al., 2011), and most grassland forbs are entomophilous. Thus, the loss of grassland plant diversity and abundance due to past forest land use can cause the loss or reduction of mutualistic pollinators in a given area (Ebeling et al., 2008; Ushimaru et al., 2018), which in turn impedes the reproductive success of entomophilous species by resulting in pollen limitation. The possible long-lasting negative impacts of this plant-pollinator feedback on grassland plant communities have rarely been examined to date. Thus, examining characteristics of plant-pollinator interactions and the extent of pollen limitation in entomophilous plants within communities should provide important insights into legacy effects on community dynamics in secondary grassland succession following deforestation.

Network approaches are useful for describing the structures of community-wide plant-pollinator interactions and predicting the dynamics of pollination communities and plant reproductive success therein (Kaiser-Bunbury et al., 2014; Magrach et al., 2021). Decreased plant and pollinator diversity may degrade pollination functions through network generalization. Low plant diversity may facilitate pollinators to increase pollinator niche overlap by limiting the diet variety (Gomez-Martinez et al., 2022). On the other hand, reduced pollinator richness may facilitate the establishment of super-generalist pollinators by reducing interspecific competition (Olesen et al., 2002; Traveset et al., 2016). Thus, both decreased plant and pollinator diversity would result in network-level generalization (Traveset et al., 2016; Gomez-Martinez et al., 2022). Highly generalized networks, in turn, may have detrimental effects on pollination function by reducing conspecific pollen receipt and promoting heterospecific pollen deposition on stigmas (Aizen & Feinsinger, 2003; Armbruster, 2006). Furthermore, reduced pollinator abundance may cause pollinator-limited reproduction in flowering plant communities. Thus, we predicted that low plant diversity would reduce pollinator richness and abundance, and cause network generalization, resulting in low pollination services and seed reproduction for entomophilous plants. Despite the advancement of such predictions in theory, there were no empirical evidence.

To test this prediction, we examined quantitative plant-pollinator networks in grasslands within ski slopes. These grasslands were established after forest removal following temporal forestation, and we use it as examples of restored grasslands (Roland et al., 2007; Inoue et al. 2021). We examined 30 plant-pollinator networks in 10 grasslands that cover a wide range of years since grassland recovery (2-84 years) and 5 reference old-growth grasslands (at least 165 years old). This unique semi-experimental study system is unparalleled in other studies, allowing us to observe how pollination networks change in the long-term dynamics of grassland recovery. Using this system, we also compared the pollination success of 15 native plant species and pollen limitation in seed set of 5 native plant species between grassland types (old and restored grasslands) and among restored grasslands with different years since forests were removed. We further examined changes in these network features and plant reproduction parameters over the years after forest cutting in the restored grasslands. We aimed to address the following questions: (1) Are plant and pollinator richness and abundance lower in restored grasslands than in old grasslands? (2) Are pollination networks in restored grasslands more generalized than in old grasslands? (3) Do more generalized networks in restored grasslands cause lower pollination success and higher pollen limited seed production in entomophilous species than do more specialized networks in old grasslands? (4) How do plant and pollinator diversity, network structure, and pollination success in communities recover with increasing time after forest cutting in restored grasslands? Based on our results, we answered these questions and discussed long-lasting legacy effects of plant-pollinator feedback.

## MATERIALS & METHODS

### Study area and plots

We examined pollination communities on 15 ski grasslands (5 old and 10 restored grasslands) at three ski resorts in the Sugadaira Highland in Nagano Prefecture, center of Japan, six slopes at Davos, seven slopes at Taro, and two slopes at Omatsu (Fig. 1, 36°51’–54’N, 138°31’–36’E, approximately 1,300–1,500 m a.s.l.) in 2021–2022. In year 1855, semi-natural grasslands (mainly pastures), which were widely maintained by grazing, burning, and mowing, covered almost the entire study area (Inoue et al., 2021). Some pastures have been used as ski slopes since the 1920s, whereas others were afforested by abandonment or plantation in the 1910s. Subsequently, from 1937, some forested areas were deforested again to create ski slopes by clearing forests. We defined the former slopes as old grasslands and the latter as restored grasslands. Note that grassland communities on restored grasslands had once disappeared during forest periods (Inoue et al., 2021). Both grassland types have usually been maintained by annual mowing in autumn, which is the conventional ski slope management (Yaida et al., 2019). We confirmed that only a single restored grassland was mowed twice a year. We established a 1000 m² (5 × 200 m) belt plot for each of the 5 old and 10 restored grasslands in the study area for both 2021 and 2022 (Table S1).

**Fig. 1.**
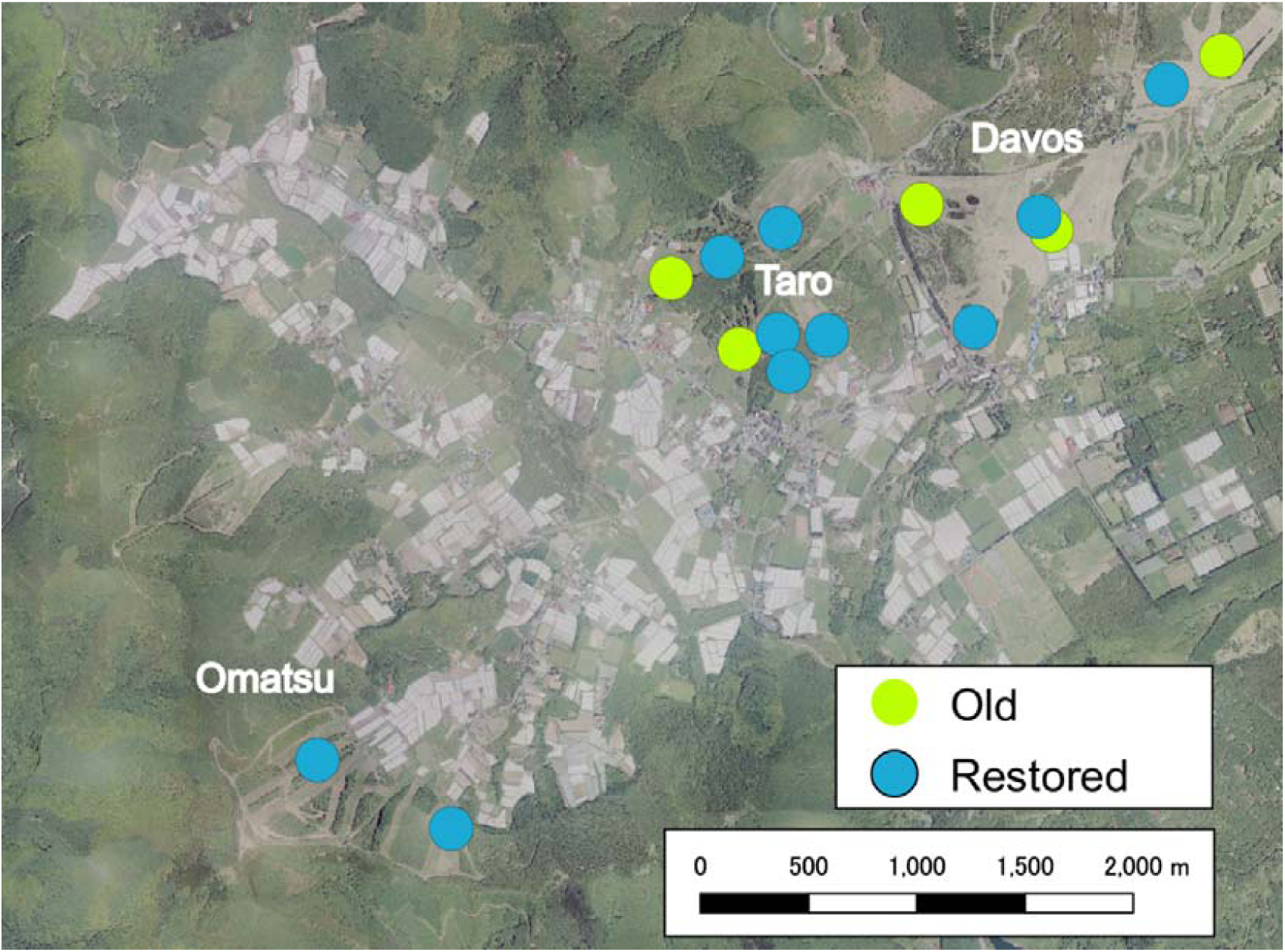
Map of the 15 study grasslands at the three ski resorts: Davos, Taro, and Omatsu in the Sugadaira Highland, Nagano Prefecture, Japan. 5 reference old (green circles) grasslands and 10 restored grasslands (blue circles) were surveyed.

### Flower-pollinator networks

Five seasonal surveys were conducted in each of 15 plots, approximately once a month, on sunny and warm days from May to September in 2021 and 2022. Mowing timing differed slightly among the grasslands, from late August to early September, respectively (Table S1). During each seasonal survey, we recorded all plant species with insect-pollinated flowers and counted the number of flowers in each plot. We then conducted a survey wherein a single person walked along each belt plot for approximately 45 min and collected flower visitors who touched the stigmas and anthers (hereafter ‘pollinators’). Each survey comprised four observations (two in the morning, 08:00–12:00 h and two in the afternoon, 12:00–16:00 h). Specimens were kept in our laboratory and identified to the species or morphospecies level. We conducted 292 and 296 observations in 2021 and 2022, respectively. In total, we constructed 30 plant-pollinator networks (15 networks×2 years). We recorded 11,413 plant–pollinator interactions between 78 flowering plants and 300 pollinator species in 2021, and 20,594 interactions between 89 flowering plants and 300 pollinator species in 2022.

To estimate the flower abundance of each species at each site and during each survey, we measured several flower parameters to calculate the visual (attraction) size (mm²) of the three sampled flowers for each plant species, following Hiraiwa and Ushimaru (2017). For each site and survey, flower abundance of each species was calculated as the total visual size of the flowers (mean visual size multiplied by the number of flowers). The number of observed individuals was used as the abundance of each pollinator species in each survey. We summarized the annual data collected from each site as a unit to calculate the network parameters for subsequent analyses.

### Network metrics

To describe the structure of each network, we calculated the following seven network metrics that are often used to examine network-level generalization and are potentially influenced by plant and pollinator richness (Traveset et al., 2016) using the bipartite package of R. (R Development Core Team). The connectance (C) reflects the proportion of possible links in a network. Complementary specialization (HL’) describes the degree of specialization between plants and pollinators in the network. Interaction strength asymmetry (ISA) describes the differences between the interaction strength of each animal species with each plant species, and vice versa, where positive values indicate a high dependence of animals on plant species, and negative values indicate the opposite. Niche overlap of flowers and pollinators (NOF and NOP, respectively) reflect the mean similarity in interaction patterns between species of the same trophic level. The weighted nestedness (WNODF) quantitatively reflects the degree of network nestedness, indicating that specialized species are preferentially associated with generalist species. Further, modularity (M) refers to the existence of subsets (modules) of closely interacting species with relatively few or no interactions with other subsets.

Because calculations of network metrics are strongly influenced by sampling biases (Blüthgen, 2010; Dormann, 2017), we corrected each metric value by calculating the z-score using null model expectation (Dormann & Strauss, 2014) as follows: z-score = (NP_obs_ – NP_null_) /σ_null_, where NP_obs_ and NP_null_ are the observed value and the mean of 10,000 null models, respectively and σ_null_ is the standard deviation calculated from the null models. To meet the null model expectations, we shuffled the entries of the observed plant-pollinator matrix of each network while maintaining the number of interactions and plant and pollinator richness; moreover, we calculated each metric for each randomisation of individual interactions (10,000 randomizations; r2dtable in the bipartite package of R). Corrected network metrics were used in all the following analyses and discussions without mentioning ‘corrected’.

PCA was conducted on the seven network metrics of each network to reduce the number of network metrics included in the data analyses. Two primary axes explained 93.1% of the total variance (PC1: 86.5%; PC2: 6.6%). The PC1 value was negatively correlated with connectance, NOF, NOP, and WNODF, and positively correlated with interaction strength asymmetry, complementary specialization, and modularity (Fig. S2). Thus, an increase in PC1 generally indicated an increase in network-level specialization (or a decrease in network-level generalization). We used only the PC1 value in our analyses because PC1 alone accounted for most of the total variances.

### Community-wide pollination success of herb species

#### Pollen receipt

– To examine the effects of network features on community-wide pollination functioning, we investigated pollen receipt on the stigmas of 6 and 14 species in 2021 and 2022, respectively (total 15 species). Plant species that were relatively widely distributed in both old and restored grasslands (Fig. S3) and that exhibited various flowers with different phenologies, morphologies, and pollination partners, were selected. During the three seasonal surveys (late June, late July, and mid-August) in 2021 and four seasonal surveys (late June, late July, mid-August, and early September) in 2022, we collected a single stigma from an open-pollinated, withered flower from each of the 20 individuals (or more than five individuals if there were <20) for each species at each site in each survey period.

Pollen receipt (number of conspecific (Con) or heterospecific (Het) pollen grains deposited on each sampled stigma) of each flower was examined under a microscope. The Con and Het pollen receipts on each stigma were standardized for each species across all study sites using *z*-scores, that is, (observed value – mean value) / standard deviation for the species, prior to subsequent analyses, thus, permitting inter-species comparison.

#### Seed set

– To examine reproductive success and pollen limitation in entomophilous plants, we investigated the seed set of 5 species out of 15 species whose pollen receipt was examined mainly in the Taro and Davos areas in mid-June, mid-July, mid-August, and early September 2022 (Fig. S3). To test pollen limitation, we conducted experimental pollination treatments on the flowers of each species at each sampling site. We marked the flowers (or inflorescences) and assigned them to two treatment groups: naturally pollinated flowers (natural flowers) and flowers with supplemental (hand) pollination (supplemental flowers). For each treatment, 15–20 individuals (or more than 5 individuals if there were <15) were examined for each species and site. Approximately 1 month after the experiment, we collected fruits and seeds from the natural and supplemental flowers of each species and dried them in our laboratory. Sites with individuals of target species <5 within and around the study plot were not investigated. In total, we collected fruits and seeds from 1,115 plant individuals.

We counted the number of mature and immature seeds and calculated the seed set (number of mature seeds/total number of seeds) for each individual plant. Furthermore, we calculated the pollen limitation index (PLI), which describes the proportional decline in seed size of naturally pollinated plants compared to supplementally pollinated plants, and is quantified as

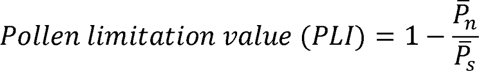

where *p̅_n_* and *p̅_s_* are the average seed sets of the natural and supplemental flowers of each species at each site, respectively. Seed set and PLI was standardized for each species across sites using z-scores prior to subsequent analyses.

### Statistical analysis

#### Flower and pollinator diversity

– We first compared the basic network parameters (total, native, and exotic flowering plant richness (i.e. number of flowering plant species) and abundance, and pollinator richness (i.e. number of pollinator species) and abundance between the old and restored grasslands. A generalized linear mixed model (GLMM), Poisson errors, and log-link function were used for count data, and Gaussian errors and identity link function were used for continuous data for this and subsequent analyses. In each model, each network parameter was included as the response variable with the grassland type (old/restored) being the explanatory variable. The effects of spatial autocorrelation on statistical analyses were considered because the study sites were not randomly distributed within the study area. Therefore, we calculated the spatial auto-covariates from the latitude and longitude measurements of all sites and incorporated the values into each GLMM (Dormann et al., 2007). Furthermore, the study year and survey time (four or five) were included as independent random terms in each model to avoid the effects of temporal pseudo-replication and sampling efforts. We did not include site as a random factor because the spatial auto-covariates already control for the differences between sites.

We also examined the relationships between these network parameters and years since grassland restoration (grassland age) began at the restored sites. We constructed GLMMs with the same response variables and random terms, including grassland age as an explanatory variable.

#### Network metrics

– To compare the network metric PC1 between the old and restored grasslands, a GLMM was constructed with network PC1 and grassland type as the response and explanatory variables, respectively. We then examined the relationship between network PC1 and grassland age in the restored grasslands using GLMMs. In each GLMM, we incorporated spatial auto-covariates as explanatory variables, and study year and survey time as independent random terms in the model.

We further examined the relationships between network PC1 and plant and pollinator diversity parameters (plant and pollinator richness and abundance) using GLMMs. Before each analysis, we calculated the variance inflation factors for each explanatory variable of the models to avoid multi-collinearity and removed variables with a variance inflation factor value of >3 (Zuur et al., 2010). In the final model, plant richness and abundance, pollinator richness, and spatial auto-covariates of the response variable were included as explanatory variables. In each GLMM, the study year and survey time were used as independent random terms.

### Plant reproduction

#### Community-wide pollination function

– We compared community-wide pollen receipt between old and restored grasslands and examined their relationship with grassland age using GLMMs. In the GLMMs, the standardised Con or Het pollen numbers on each stigma for each species and grassland type or age were included as the response and explanatory variables, respectively. Furthermore, we constructed a GLMM to examine the relationship between pollen receipt and network PC1, in which the response variable was the standardized Con or Het pollen receipt for each stigma, and the explanatory variable was PC1. To examine the potential effect of pollinator visitation on pollen receipt, we incorporated the visiting frequency (visits per flower area (cm²)) for each species at each site as a covariate. Prior to analysis, visiting frequency was standardized to permit inter-species comparisons. The spatial auto-covariate of the response variable was also included as a covariate. Study year and survey identities were included in the models as nested random terms, and plant family and species identities were also fitted as nested random terms to account for phylogenetic constraints.

#### Seed set and pollen limitation

– We compared seed set and PLI between old and restored grasslands. First, we examined a GLMM wherein the standardized seed set was a response variable, and four flower groups (2 treatments × 2 grassland types) were explanatory variables, whereas the spatial auto-covariate was included as a covariate and survey identity and plant family and species identities were independent and nested random terms. Subsequently, we conducted a multiple comparison among flower groups by Tukey’s method using the ‘multcomp’ package (Hothorn, Bretz, & Westfall, 2008).

Next, we examined the GLMM with PLI and grassland type as the response and explanatory variables, respectively. The survey identity was included as an independent random variable in the model. Plant family and species identities were also fitted as nested random terms to account for phylogenetic constraints.

The significance of each explanatory variable in the GLMMs was examined using the Wald test. All analyses were performed using R.

## RESULTS

### Richness and abundance of flowers and pollinators

Restored grassland sites had significantly lower plant species richness and their richness increased significantly with grassland age (Fig. 2a, Table S2), which mainly driven by native flowering plants (Fig. S4). Furthermore, flower abundance had no significant relationship with grassland type or age at the restored sites (Fig. 2b, Table S2). However, the flower abundances of native and exotic species significantly increased and decreased with grassland age in the restored sites, respectively (Fig. S4). Meanwhile. pollinator richness was similar among slope types and was not significantly correlated with grassland age at the restored sites, whereas pollinator abundance was significantly higher at the old sites than at the restored sites (Fig. 2c, d, Table S2).

**Fig. 2.**
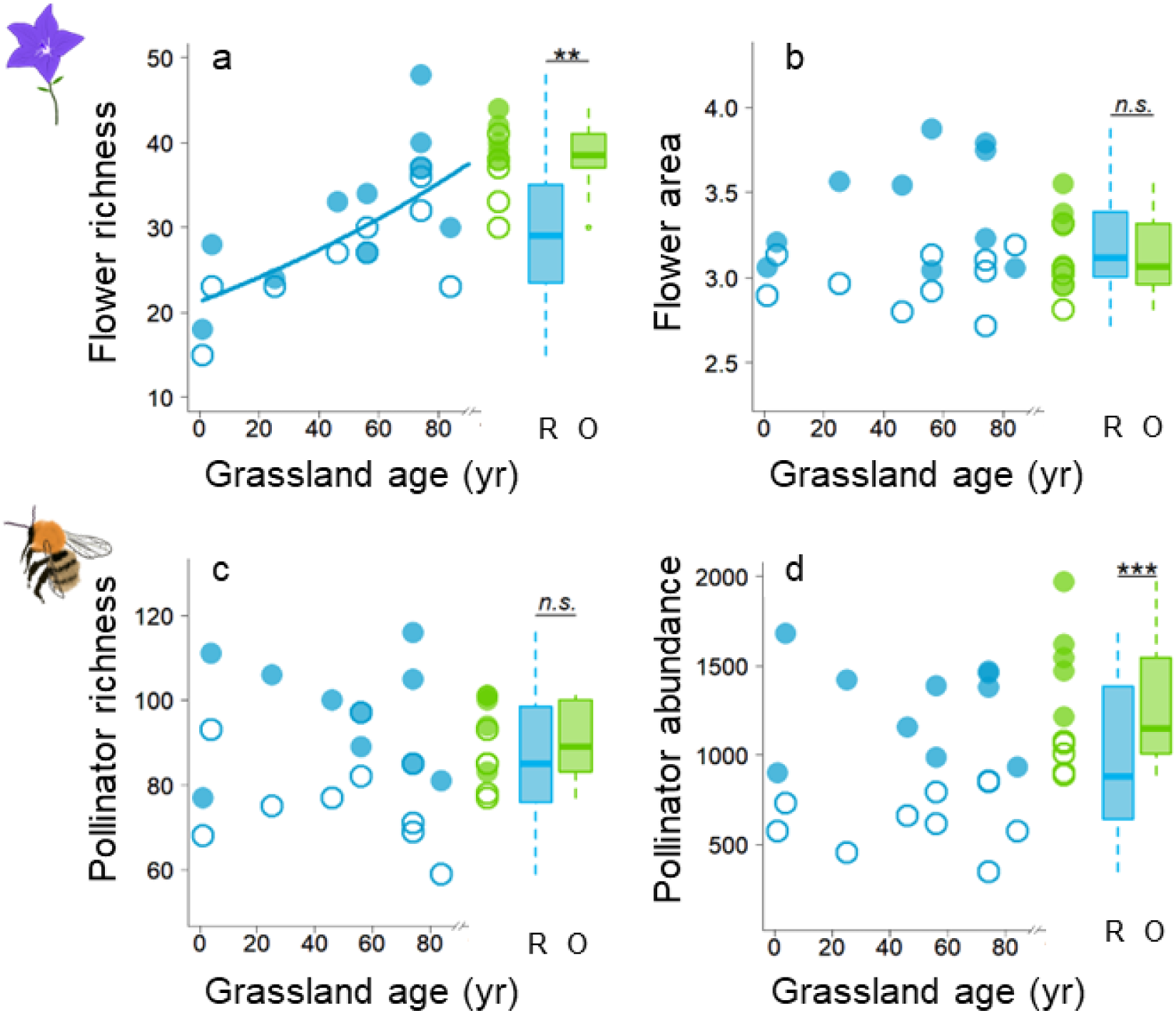
Comparison of flowering plant richness and flower area (log10(cm²)), and pollinator richness and abundance examined in 2021 (open circle) and 2022 (closed circle) between old grasslands (green, O) and restored grasslands (blue, R), and their relationships with grassland age (year) in the restored grasslands. The solid lines represent significant regressions from the generalized linear mixed models (GLMMs) (Table S2). Boxplots indicate plant diversity metrics for O and R plots; central bars indicate the median, and asterisks indicate significant differences (***, *p < 0.001*; **, *p < 0.01*; n.s., *p > 0.05*).

### Network metrics and pollination success

Network PC1, which indicates the degree of network-level generalization, was significantly higher at the restored sites than at the old sites, and significantly decreased with grassland age (Fig. 3a, Table S2). Furthermore, the total flowering plant richness had a significant positive relationship with PC1 (Fig. 3b, Table S2). Community-wide Con pollen receipt by plants at the restored sites was significantly lower than that at old sites and significantly increased with grassland age at the restored sites (Fig. 4a, Table S2). Community-wide Het pollen receipt in restored sites was higher than that at the old sites and significantly decreased with grassland age (Fig. 4b, Table S2). Network PC1 had significant positive effects on community-wide Con pollen receipt (Fig. 4c, d, Table S2).

**Fig. 3.**
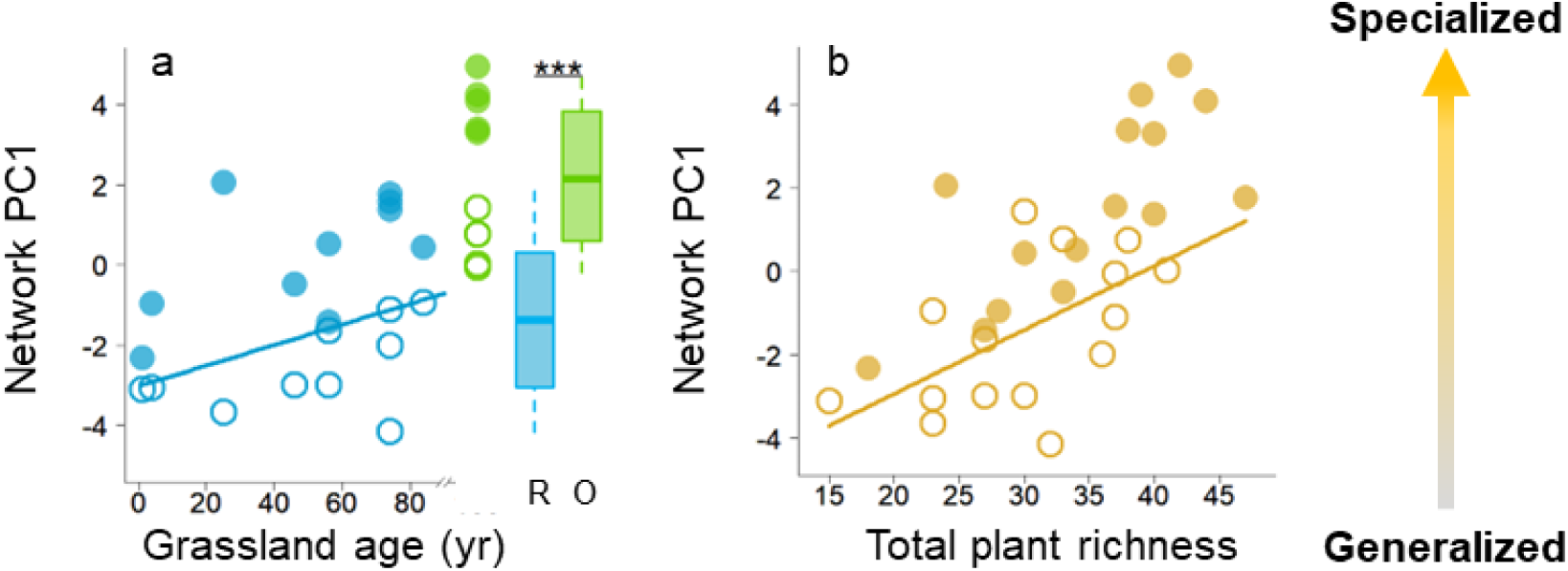
(a) Comparison of network PC1 examined in 2021 (open circle) and 2022 (closed circle) between old grasslands (green, O) and restored grasslands (blue, R), and their relationships with grassland age (year) in restored grasslands. (b) Relationships of network PC1 with plant richness. The solid lines represent significant regressions from the GLMMs (Table S2). Boxplots indicate network PC1 values for R and O plots; central bars indicate the median, and asterisks indicate significant differences (***, *p < 0.001*).

**Fig. 4.**
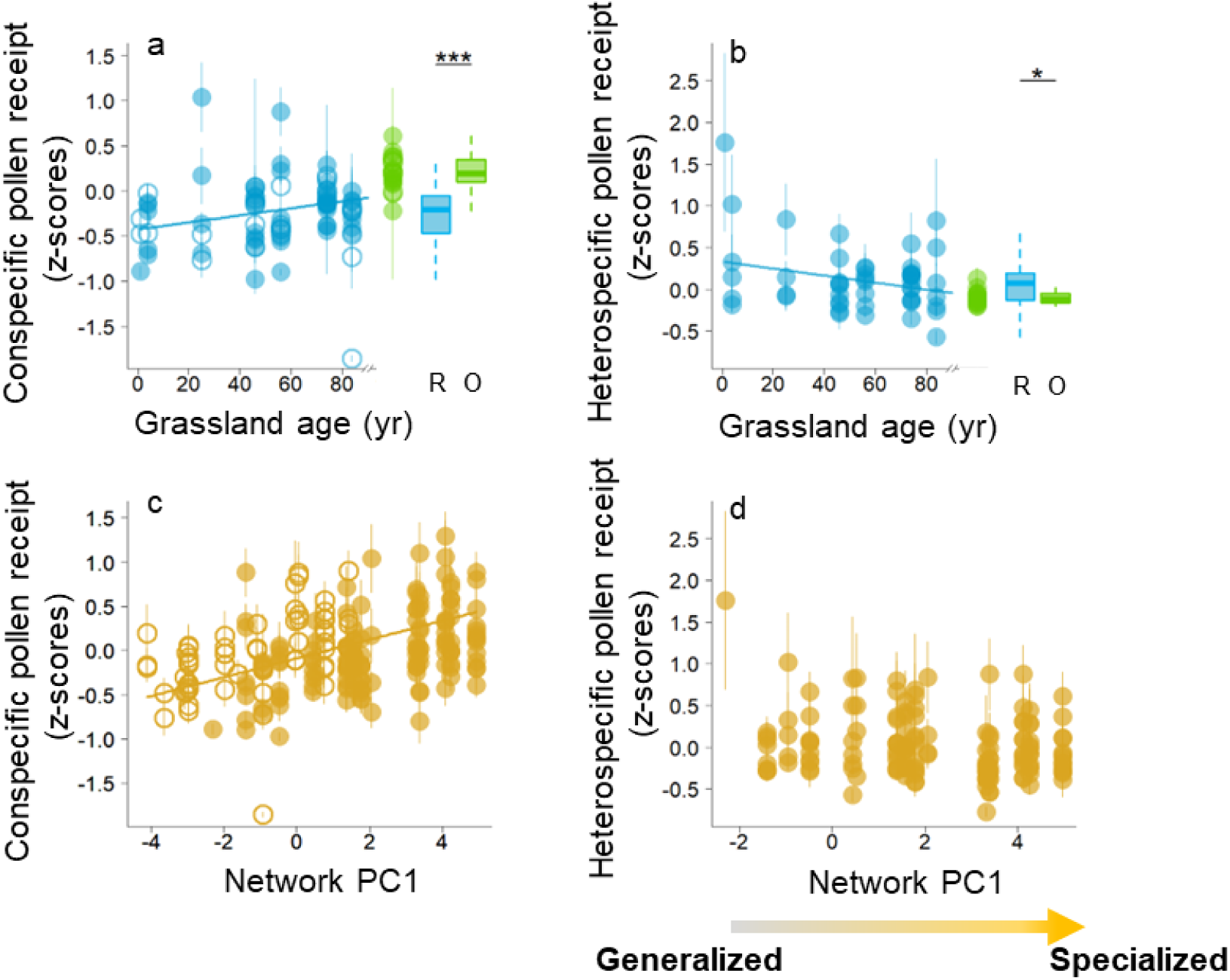
(a, c) Comparison of conspecific (Con) and heterospecific (Het) pollen receipt (z-scores) examined in 2021 (open circle) and 2022 (closed circle) between old grasslands (green, O) and restored grasslands (blue, R), and their relationships with grassland age (year) in restored grasslands. (b, d) Relationships of these pollen receipt values with network PC1, respectively. The solid lines represent significant regressions from the GLMMs (Table S2). Boxplots indicate pollen receipt values for R and O plots; central bars indicate the median, and asterisks indicate significant differences (***, *p < 0.001*; *, *p < 0.05*).

### Seed set and pollen limitation

The seed set at the old sites did not differ between the natural and supplemental pollination treatments (Fig. 5a, Table S2), whereas the natural seed set was significantly lower than that of flowers with supplemental pollination at the restored sites (Fig. 5a, Table S2). The degree of pollen limitation was significantly higher at the restored sites than at the old sites (Fig. 5b, Table S2). Furthermore, the degree of pollen limitation was significantly negatively correlated with the amount of Con pollen received (Fig. 5c, Table S2).

**Fig. 5.**
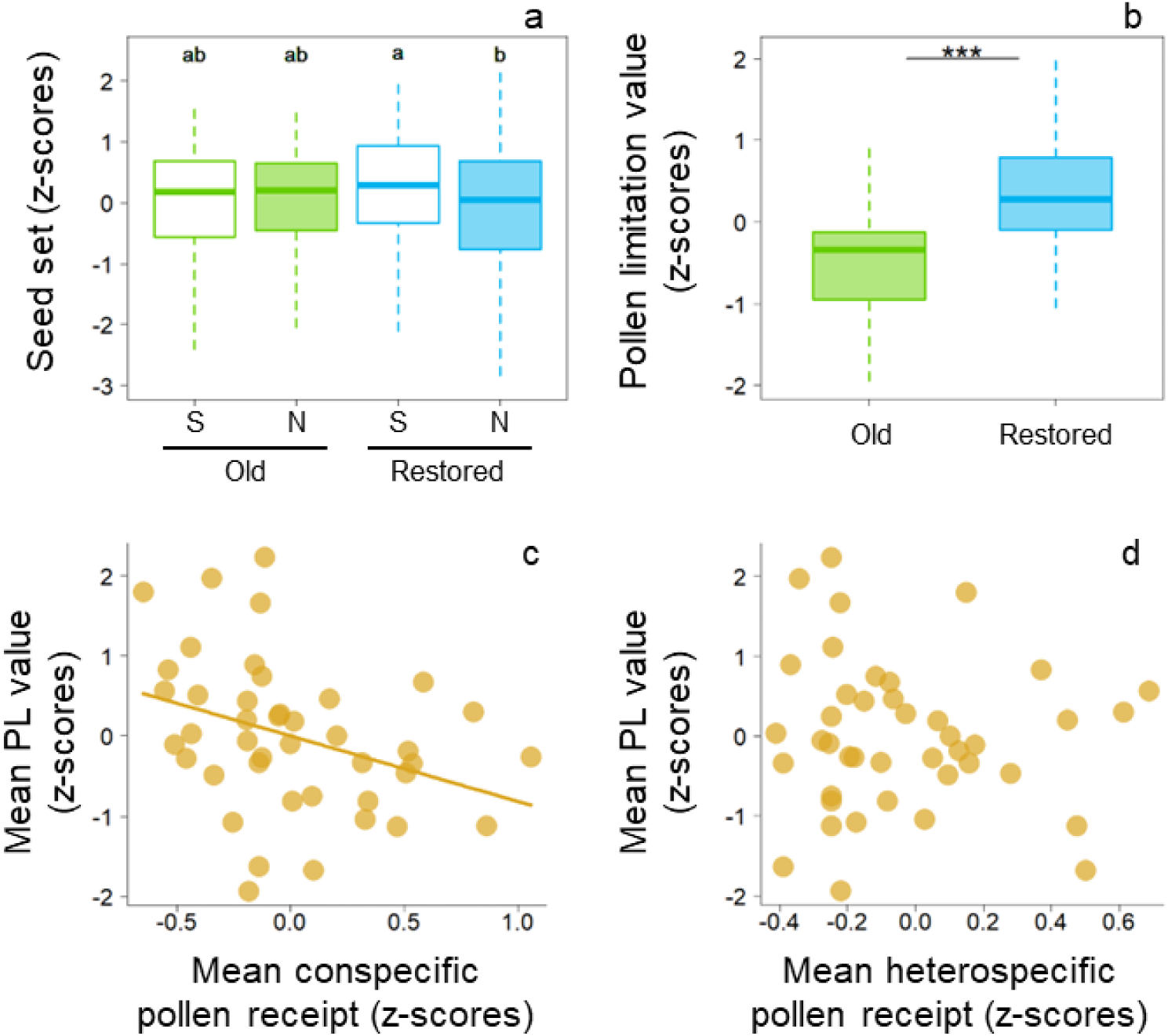
(a) Seed set (z-scores) of supplemental (open, S) and natural (closed, N) pollinated plants in old (O) – and restored (R) grasslands. Different letters indicate significant (*p < 0.05*) differences among treatments based on Tukey’s method (Table S2). (b) Comparison of pollen limitation (z-scores) between grassland types. Boxplots represent medians (central bars), and first and third quartiles (box perimeters). Asterisks indicate significant differences (***, *p < 0.001*). (c, d) Relationships of mean pollen limitation values (z-scores) with mean Con and Het pollen receipt (z scores) for each species of each site. The solid lines represent significant regressions from the GLMMs (Table S2).

## DISCUSSION

Our results showed that native plant richness, the degree of network-level specialization, and pollination and reproduction success of native plants were significantly diminished by past forestation at the restored grasslands. Moreover, we found that it can take more than 80 years after the re-introduction of grassland management to recover pollination communities and their function. Additionally, we demonstrated that the high pollen limitation of native plants was caused by network generalization via flower-poor communities in the restored grasslands. Thus, the negative impacts of initially low plant diversity may hinder grassland plant restoration. To our knowledge, this is the first study to highlight the legacy effects of past forestation on grassland plant-pollinator networks and pollination functions. The role of this long-lasting legacy effect on the slow recovery process of ski grassland restoration has been discussed comprehensively below.

Compared with the old grasslands, restored grasslands had a lower diversity of native entomophilous flowers, as reported previously (Inoue et al., 2021; Yaida et al., 2022). Furthermore, we observed the dominance of a few exotic species immediately after tree removal and the gradual reduction with the recovery of native flowers (Fig. S4). Thus, restored grasslands after forest clear-cutting may experience a succession of flower communities from those dominated by exotic species to those dominated by native species via long-term grassland restoration. The succession will take over 70-80 years to reach a similar level to that of old grasslands. Unexpectedly, pollinator richness did not differ significantly between grassland types and was not influenced by the grassland management period despite the low richness of flowering plants (Fig. 2a). Pollinators are highly mobile insects that track resources actively, and therefore, they can move from surrounding habitats more easily, buffering differences between grasslands.

As expected, plant-pollinator networks were more generalized in the restored grasslands with lower plant richness than in the old grasslands (Fig. 3a). In previous studies, generalized networks were typically found in communities with low pollinator richness or low richness of both plants and pollinators (Olesen & Jordan, 2002; Traveset et al., 2016; Tommasi et al., 2021; Gomez-Martinez et al., 2022). Our study firstly reports that network-level generalization is caused solely by low plant richness, independent of pollinator richness. Low plant richness resulted in more generalized networks, likely by promoting pollinator niche overlap, owing to limited flowering species resources. Moreover, the mass flowering of several invasive plants may facilitate the concentrated foraging of diverse pollinators by these species (Vila et al., 2009). Generally, to reduce interspecific competition and increase their foraging efficiency despite their wide diet breadth, pollinator species compete against each other and segregated their niches when flower resources are taxonomically and functionally diverse (Frund et al., 2013; Hiraiwa & Ushimaru, 2017, 2024). Thus, an increase in plant richness over time after resuming mowing management decreased network-level generalization (i.e. network-level specialization increased) in restored grasslands (Fig. 3b).

Our findings support the hypothesized degradation of community-wide pollination functions (lower conspecific and higher heterospecific pollen receipt of native plants) in generalized networks of restored grasslands (Fig. 4a, b). Generalized networks can enhance interspecific flower movements of pollinators, which likely reduces conspecific pollen transfer and increases heterospecific pollen deposition (Aizen & Feinsinger, 2003; Armbruster, 2006; Hiraiwa & Ushimaru, 2024). In our study, the degree of network generalization had a significant negative effect on conspecific pollen flow (Fig. 4c, d). Notably, the stigmas received much more conspecific pollen than heterospecific pollen in all studied species (Fig. S5). Thus, generalized networks significantly decreased conspecific pollen deposition on stigmas rather than increasing heterospecific pollen deposition. Furthermore, the pollination function in the restored grasslands improved during the grassland management period by increasing network specialization.

We observed a more pollen-limited seed set of native plant species in the restored grasslands than in the old grasslands (Fig. 5a). Moreover, pollen limitation was caused by low conspecific pollen receipt (Fig. 5c), which was consistent with previous findings (Waser & Fugate, 1986; Flanagan et al., 2009). Thus, low conspecific pollen deposition due to network generalization was the main factor causing the decline in seed reproduction in the restored grasslands. Low seed reproduction in native plants via changes in pollinator communities and network structures resulting from low plant diversity limits the recovery of native species in restored grasslands (Suding et al., 2004). Thus, we found a negative feedback effect starting from low plant richness in younger restored grasslands. Interspecific pollinator movement can reduce seed production not only through conspecific pollen loss but also through enhanced heterospecific pollen deposition (Wilcock & Neiland, 2002; Morales & Traveset, 2008; Flanagan et al., 2009). The effects of heterospecific pollen deposition on plant fitness depend on species diversity and identity, and the quantity of heterospecific pollen (Arceo-Gomez and Ashman, 2011). Our results showed that heterospecific pollen deposition had no significant effect on pollen limitation. However, we examined pollen limitation in the seed set of only five species. Thus, further research is warranted to investigate the relationship between pollen transfer and reproductive success in more species to further generalise our findings. Contrary to our findings, Kaiser-Bunbury et al. (2017) reported that ecological restoration by removal of exotic plants in Inselberg plant communities facilitated plant-pollinator network generalization, which improved the pollination function for native plants. Network structure was found to be a suitable indicator of pollination function, but there are still few examples, and no consistent evidence is available. Thus, future studies should focus on the relationships between the network structure and pollination functions in various ecosystems.

This study was the first to provide evidence of the long-lasting legacy effects of past forestation on pollination networks and functions via the negative impacts of low plant richness feedback through changing network properties in restored grasslands. Low plant richness caused generalized plant-pollinator networks, which in turn resulted in low pollination function and high pollen limitation in restored grasslands with a short management history. Insect-pollinated plants are known to disappear rapidly after grassland abandonment (Sõber et al., 2023) and also slower to recover after grassland restoration (Alberht et al., 2021). The slow recovery of insect-pollinated plants may be caused by network generalization due to initial low plant diversity after forest cutting. Even if a given plant species has successfully recovered in terms of seed dispersal and establishment, it is unlikely to reproduce effectively without the restoration of highly specialized plant-pollinator networks which is supported by high plant richness. These findings suggest that the recovery of specialized plant-pollinator networks by enhancing plant diversity is essential for restoring pollination function. For quicker grassland restoration, it may be effective to facilitate the establishment of highly specialized pollination networks by seeding or planting diverse native plants collected from neighbouring areas while avoiding genetic contamination. We emphasize the importance of the entire mutualistic network structure for the restoration of grassland plant communities. Overall, pollination network monitoring appears as a useful approach for checking the soundness of restored ecosystems and is recommended for restoring plant communities in various ecosystems with high insect-pollinated plant richness.

## Acknowledgement

We thank the land owners of our study sites, which are ‘Sugadaira Bokujo’, ‘Hatsune-kan’, ‘Imai-kan’, and ‘Jozan-kan’; and the management companies, ‘Sugadaira Pine beak ski’, ‘Sugadaira Ski House Co., Ltd.’, ‘Oku Davos Snow Park’, and ‘HARE Sugadaira-Kogen Snow Resort’ for permitting to conduct field surveys. We greatly thank Carlos Martinez-Nuñez for valuable comments. This work was supported by the Sasakawa Scientific Research Grant from The Japan Science Society, the fund of Nagano Prefecture to promote scientific activity to G.S.H., and a GrantLinLAid for Scientific Research Programs (KAKENHI no. 19H03303) from the Japan Society for the Promotion of Science to A.U.

## Author contribution

G.S.H. and A.U. conceived the ideas and designed methodology; G.S.H. basically collected and analysed the data; H.S.I and A.U collected flower trait data; T.I and T.K examined the history of our study grassland; G.S.H. wrote first draft together with A.U. All authors contributed critically to the drafts and gave final approval for publication.

