## Supplement1 for "Long-lasting negative effects of past forestation on grassland pollination networks and their function"

**Supplemental materials**

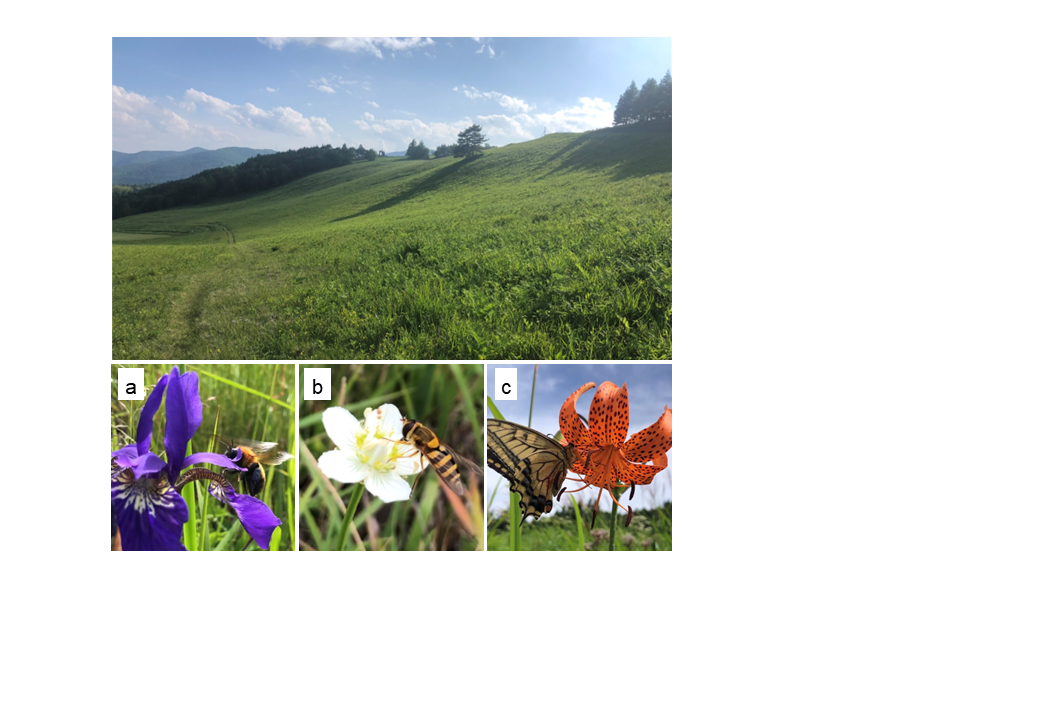

**Fig. S1** Photos of the studied grasslands and pollinators of main taxonomic groups. (a) *Bombus diversus diversus* visiting *Iris sanguinea*, (b) *Syrphus torvus* visiting *Parnassia palustris*, (c) *Papilio machaon* foraging on *Lilium leichtlinii*.

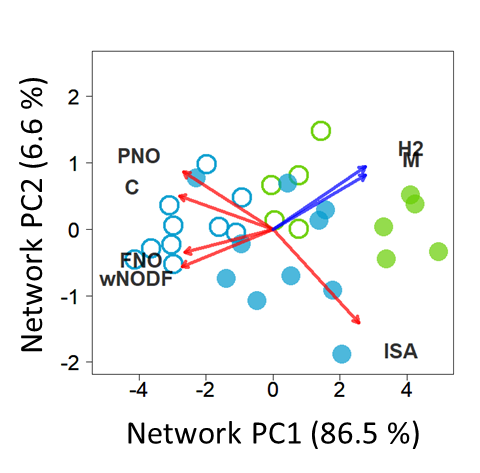
**Fig. S2** PCA bi-plot for seven network variables (calculated from connectance; weighted NODF; flower and pollinator niche overlap, H2, modularity and ISA) in 2021 (open circle) and 2022 (closed circle). The first two PCA axes explain 93.1 % of the total variance (PC1: 86.5%, PC2: 6.6%). PC1 was negatively correlated with connectance, weighted NODF, and flower and pollinator niche overlap, and positively correlated with H2, modularity and ISA. PC2 was positively correlated with connectance, pollinator niche overlap, H2 and modularity, and negatively with weighted NODF, Flower niche overlap and ISA. Red lines indicate the metrics positively correlated with network generalization, and blue lines indicate the metrics positively correlated with network specialization (Traveset et al., 2016).

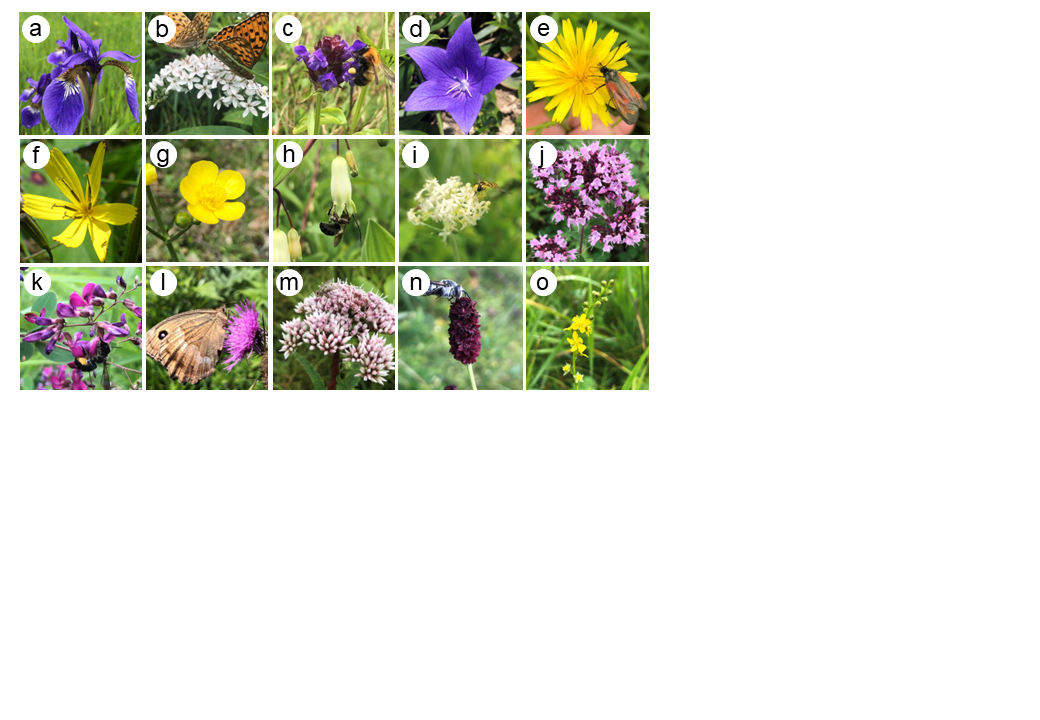

**Fig. S3** Photos of native plants examined in pollen receipt surveys. (a) *Iris sanguinea*, (b) *Lysimachia clethroides*, (c) *Prunella vulgaris*, (d) *Platycodon glandiflorus*, (e) *Picris hieracioides*, (f) *Ixeridium dentatum*, (g) *Ranunculus japonicus*, (h) *Polygonatum dentatum*, (i) *Galium verum*, (j) *Thymus quinquecostatus*, (k) *Lespedeza bicolor*, (l) *Cirsium oligophyllum*, (m) *Eupatorium glehnii*, (n) *Sanguisorba officinalis*, and (o) *Agrimonia pilosa*.

The species (a)–(e) were examined in both pollen receipt and seed set surveys.

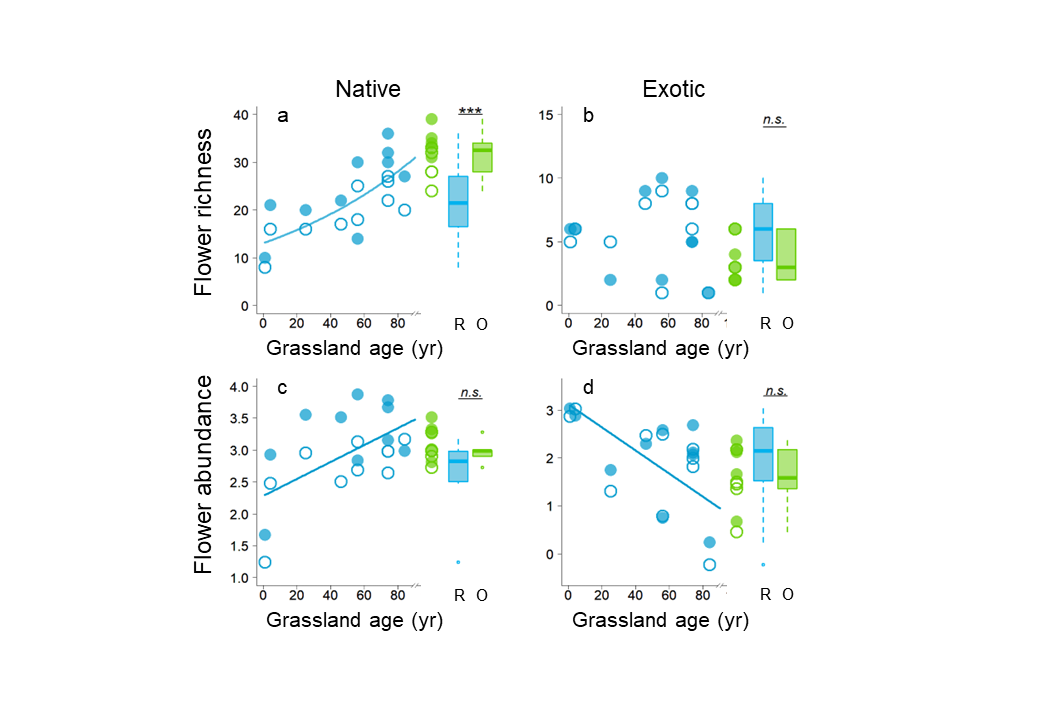
**Fig. S4** Comparison of richness and flower area (log10(cm²)) of native and exotic plants examined in 2021 (open circle) and 2022 (closed circle) between old grasslands (green, O) and restored grasslands (blue, R), and their relationships with grassland age (year) in the restored grasslands. The solid lines represent significant regressions from the generalized linear mixed models (GLMMs) (Table 1). Boxplots indicate plant diversity metrics for O and R plots; central bars indicate the median, and asterisks indicate significant differences (***, *p < 0.001*; **, *p < 0.01*; n.s., *p > 0.05*).

**Fig. S5** Comparison of the number of pollen grains (mean ± *SE*) with pollen t
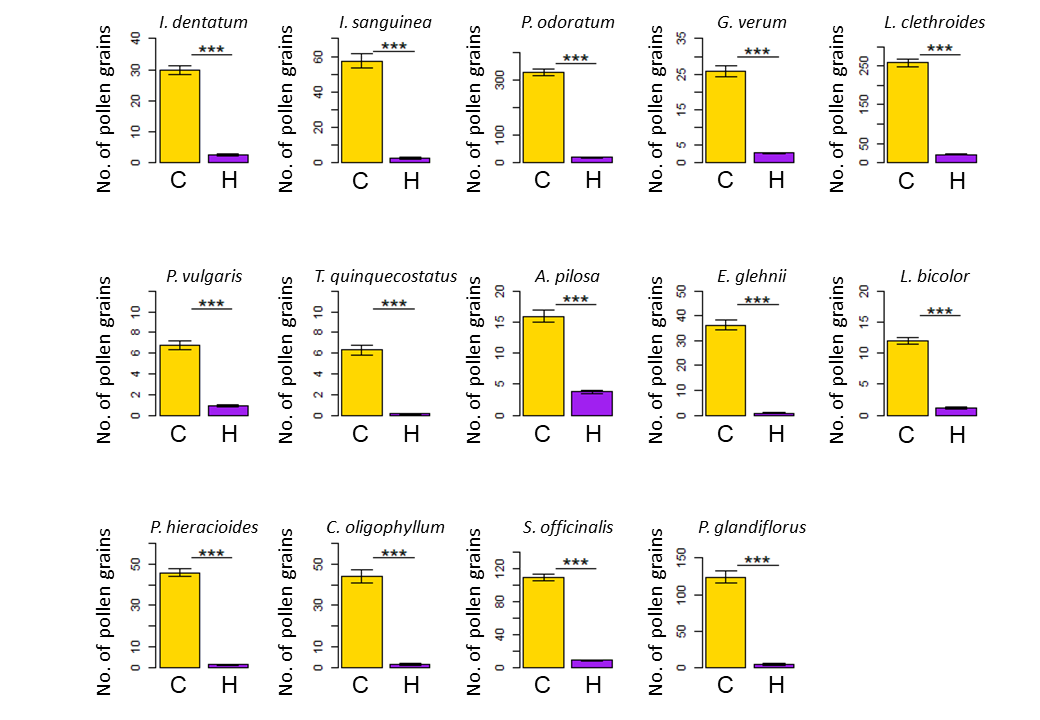
ype: conspecific (C) or heterospecific (F) for each species. Asterisks indicate significant differences (***, *p < 0.001*).

Table S1 Basic information of the study site.

| **Site** | **Ski area** | **Grassland type** | **Longitude** | **Latitude** | **Deforestation** | **Survey period  (2021)** | **Survey period  (2022)** |
| --- | --- | --- | --- | --- | --- | --- | --- |
| Hinode(n) | Taro | Restored | 138.3345 | 36.533 | 1965 | May-September | May-August |
| Hinode(o) | Taro | Restored | 138.335 | 36.5321 | 1947 | May-August | May-September |
| Tengu | Taro | Restored | 138.3365 | 36.5333 | 1947 | May-September | May-September |
| Omotetarou | Taro | Old | 138.3318 | 36.5324 | NA | May-September | May-September |
| Uratarou | Taro | Restored | 138.3348 | 36.5381 | 2017 | May-September | May-September |
| Shirogane(n) | Taro | Restored | 138.3313 | 36.5367 | 1975 | May-September | May-September |
| Shirogane(o) | Taro | Old | 138.3287 | 36.5362 | NA | May-September | May-September |
| Okudabosu | Davos | Old | 138.3583 | 36.5447 | NA | May-September | May-September |
| Nishidabosu | Davos | Restored | 138.3547 | 36.5433 | 1937 | May-September | May-September |
| Omotedabosu | Davos | Old | 138.3485 | 36.5374 | NA | May-September | May-September |
| Uradabosu | Davos | Old | 138.342 | 36.5385 | NA | May-September | May-September |
| Schneider | Davos | Restored | 138.3444 | 36.5335 | 1947 | May-September | May-September |
| Slope | Davos | Restored | 138.3479 | 36.538 | 2020 | May-September | May-September |
| Omatsu(g) | Omatsu | Restored | 138.3163 | 36.5123 | 1965 | May-September | May-September |
| Omatsu(w) | Omatsu | Restored | 138.3097 | 36.5164 | 1996 | May-August | May-September |

**Table S2** Results of generalized linear mixed model (GLMM) analysis. Old grassland is used as a baseline for the variable, grassland type in the models. Significant effects are indicated in bold (*P < 0.05*).

| Response variables | Explanatory variable | Coefficients | | *z* | *P* |
| --- | --- | --- | --- | --- | --- |
|  |  | Estimate | SE |  |  |
| Total plant richness | **Grassland type (Restored)** | -0.244 | 0.082 | -2.974 | **<0.01** |
|  | Spatial autocovariate | -6.24E-06 | 2.61E-05 | -0.239 | 0.811 |
|  | **Intercept** | 3.657 | 0.094 | 38.942 | **<0.001** |
| Analysis in Restored grasslands | |  |  |  |  |
|  | **Grassland age** | 0.006 | 0.002 | 3.635 | **<0.001** |
|  | Spatial autocovariate | 1.18E-05 | 2.27E-05 | 0.518 | 0.604 |
|  | **Intercept** | 3.029 | 0.104 | 29.069 | **<0.001** |
| Total flower area | Grassland type (Restored) | 0.073 | 0.086 | 0.846 | 0.405 |
|  | Spatial autocovariate | -1.47E-04 | 1.18E-04 | -1.252 | 0.222 |
|  | **Intercept** | 3.138 | 0.235 | 13.375 | **<0.01** |
| Analysis in Restored grasslands | |  |  |  |  |
|  | Grassland age | 0.002 | 0.002 | 0.911 | 0.3763 |
|  | Spatial autocovariate | -6.84E-05 | 2.16E-04 | -0.316 | 0.756 |
|  | **Intercept** | 3.076 | 0.026 | 11.709 | **<0.01** |
| Total pollinator richness | Grassland type (Restored) | -0.024 | 0.042 | -0.586 | 0.558 |
|  | Spatial autocovariate | -1.59E-07 | 2.09E-06 | -0.076 | 0.939 |
|  | Intercept | 4.480 | 0.082 | 54.608 | **<0.001** |
| Analysis in Restored grasslands | |  |  |  |  |
|  | Grassland age | -0.002 | 0.001 | -1.891 | 0.059 |
|  | **Spatial autocovariate** | 1.05E-05 | 3.22E-06 | 3.246 | **<0.01** |
|  | **Intercept** | 4.414 | 0.122 | 36.299 | **<0.001** |
| Total pollinator abundance | **Grassland type (Restored)** | -0.220 | 0.012 | -18.771 | **<0.001** |
|  | **Spatial autocovariate** | -5.74E-07 | 5.64E-08 | -10.183 | **<0.001** |
|  | **Intercept** | 6.974 | 0.275 | 25.388 | **<0.001** |
| Analysis in Restored grasslands | |  |  |  |  |
|  | Grassland age | -1.05E-04 | 2.66E-04 | -0.396 | 0.692 |
|  | **Spatial autocovariate** | 4.15E-07 | 1.04E-07 | 3.991 | **<0.001** |
|  | **Intercept** | 6.617 | 0.321 | 20.633 | **<0.001** |
| Network PC1 | **Grassland type (Restored)** | 2.947 | 0.424 | 6.952 | **<0.001** |
|  | Spatial autocovariate | -0.001 | 0.001 | -1.676 | 0.106 |
|  | Intercept | -1.196 | 1.768 | -0.676 | 0.573 |
| Analysis in Restored grasslands | |  |  |  |  |
|  | **Grassland age** | -0.026 | 0.008 | -3.204 | **<0.01** |
|  | Spatial autocovariate | 0.001 | 0.002 | 0.625 | 0.541 |
|  | Intercept | 2.828 | 1.610 | 1.757 | 0.233 |
| Network PC1 | **Plant richness** | -0.153 | 0.048 | -3.202 | **<0.01** |
|  | Pollinator richness | 0.020 | 0.029 | 0.690 | 0.497 |
|  | Plant abundance | 0.565 | 1.266 | 0.447 | 0.659 |
|  | Spatial autocovariate | -0.001 | 0.001 | -0.739 | 0.467 |
|  | Intercept | 0.133 | 8.617 | 0.015 | 0.988 |
| Conspecific pollen receipt (z score) | **Grassland type (Restored)** | -0.390 | 0.035 | -11.06 | **<0.001** |
|  | Spatial autocovariate | 8.55E-07 | 2.37E-06 | 0.36 | 0.718 |
|  | **Intercept** | 0.215 | 0.024 | 8.92 | **<0.001** |
| Analysis in Restored grasslands | |  |  |  |  |
|  | **Grassland age** | 0.004 | 0.001 | 4.11 | **<0.001** |
|  | Spatial autocovariate | -1.16E-06 | 2.39E-06 | -0.48 | 0.629 |
|  | **Intercept** | -0.434 | 0.076 | -5.72 | **<0.001** |
| Conspecific pollen receipt (z score) | **Network PC1** | -0.108 | 0.010 | -10.49 | **<0.001** |
|  | Visiting frequency per flower area (z-score) | -0.010 | 0.016 | -0.62 | 0.54 |
|  | Spatial autocovariate | -3.52E-06 | 2.61E-06 | -1.35 | 0.174 |
|  | Intercept | -0.072 | 0.122 | -0.59 | 0.55 |
| Heterospecific pollen receipt (z score) | **Grassland type (Restored)** | 0.083 | 0.042 | 2.00 | **0.0459** |
|  | **Spatial autocovariate** | -1.42E-04 | 2.41E-05 | -5.91 | **<0.001** |
|  | Intercept | -0.026 | 0.031 | -0.83 | 0.4087 |
| Analysis in Restored grasslands | |  |  |  |  |
|  | **Grassland age** | -0.004 | 0.001 | -3.04 | **<0.01** |
|  | **Spatial autocovariate** | -2.28E-04 | 9.49E-05 | -2.40 | **0.016** |
|  | **Intercept** | 0.373 | 0.095 | 3.92 | **<0.001** |
| Heterospecific pollen receipt (z score) | PC1 | 0.019 | 0.011 | 1.77 | 0.0764 |
|  | Visiting frequency per flower area (z-score) | 0.027 | 0.020 | 1.36 | 0.174 |
|  | **Spatial autocovariate** | -1.48E-04 | 2.33E-05 | -6.36 | **<0.001** |
|  | **Intercept** | 0.062 | 0.030 | 2.05 | **0.04** |
| Seed set (z score) | Supplemental in Old - Natural in Old | -0.001 | 0.088 | -0.016 | 1 |
|  | Supplemental in Old - Supplemental in Restored | -0.191 | 0.086 | -2.217 | 0.118 |
|  | Supplemental in Old - Natural in Restored | 0.129 | 0.084 | 1.531 | 0.419 |
|  | Natural in Old - Supplemental in Restored | -0.192 | 0.085 | -2.263 | 0.107 |
|  | Natural in Old - Natural in Restored | 0.128 | 0.083 | 1.537 | 0.415 |
|  | Supplemental in Restored - Natural in Restored | 0.320 | 0.080 | 3.999 | **<0.001** |
| Pollen limitation value (z score) | **Grassland type (Restored)** | 0.898 | 0.245 | 3.665 | **<0.001** |
|  | **Spatial autocovariate** | -0.013 | 0.005 | -2.700 | **0.010** |
|  | Intercept | -0.337 | 0.189 | -1.780 | 0.083 |
| Pollen limitation value  (z score) | **Conspecific pollen receipt (z score)** | -0.821 | 0.333 | -2.465 | **0.018** |
|  | Heterospecific pollen receipt (z score) | 0.127 | 0.483 | 0.262 | 0.795 |
|  | **Spatial autocovariate** | -0.013 | 0.005 | -2.471 | **0.018** |
|  | Intercept | 0.164 | 0.149 | 1.100 | 0.278 |
